# Independent mechanisms for acquired salt tolerance versus growth resumption induced by mild ethanol pretreatment in *Saccharomyces cerevisiae*

**DOI:** 10.1101/445726

**Authors:** Elizabeth A. McDaniel, Tara N. Stuecker, Manasa Veluvolu, Audrey P. Gasch, Jeffrey A. Lewis

**Author notes:** Present Address: Department of Bacteriology, University of Wisconsin–Madison, Madison, WI.

## Abstract

All living organisms must recognize and respond to various environmental stresses throughout their lifetime. In natural environments, cells frequently encounter fluctuating concentrations of different stressors that can occur in combination or sequentially. Thus, the ability to anticipate an impending stress is likely ecologically relevant. One possible mechanism for anticipating future stress is acquired stress resistance, where cells pre-exposed to a mild sub-lethal dose of stress gain the ability to survive an otherwise lethal dose of stress. We have been leveraging wild strains of *Saccharomyces cerevisiae* to investigate natural variation in the yeast ethanol stress response and its role in acquired stress resistance. Here, we report that a wild vineyard isolate possesses ethanol-induced cross-protection against severe concentrations of salt. Because this phenotype correlates with ethanol-dependent induction of the *ENA* genes, which encode sodium efflux pumps already associated with salt resistance, we hypothesized that variation in *ENA* expression was responsible for differences in acquired salt tolerance across strains. Surprisingly, we found that the *ENA* genes were completely dispensable for ethanol-induced survival of high salt concentrations in the wild vineyard strain. Instead, the *ENA* genes were necessary for the ability to resume growth on high concentrations of salt following a mild ethanol pretreatment. Surprisingly, this growth acclimation phenotype was also shared by the lab yeast strain despite lack of *ENA* induction under this condition. This study underscores that cross protection can affect both viability and growth through distinct mechanisms, both of which likely confer fitness effects that are ecologically relevant.

**IMPORTANCE:** Microbes in nature frequently experience “boom or bust” cycles of environmental stress. Thus, microbes that can anticipate the onset of stress would have an advantage. One way microbes anticipate future stress is through acquired stress resistance, where cells exposed to a mild dose of one stress gain the ability survive an otherwise lethal dose of a subsequent stress. In the budding yeast *Saccharomyces cerevisiae,* certain stressors can cross protect against high salt concentrations, though the mechanisms governing this acquisition of higher stress resistance are not well understood. In this study, we took advantage of wild yeast strains to understand the mechanism underlying ethanol-induced cross protection against high salt concentrations. We found that mild ethanol stress allows cells to resume growth on high salt, which involves a novel role for a well-studied salt transporter. Overall, this discovery highlights how leveraging natural variation can provide new insights into well-studied stress defense mechanisms.

## INTRODUCTION

All organisms experience stress and must respond to environmental perturbations throughout their lifetime. Unlike animals, unicellular organisms generally lack the ability to escape stressful environments. Thus, microbes have evolved sophisticated stress defense strategies such as genome rearrangements, small molecule synthesis, and dynamic gene expression programs to enable stress acclimation (1–3). The model eukaryote *Saccharomyces cerevisiae* responds to diverse stresses by coordinating the expression of condition-specific genes with a large, common gene expression program called the environmental stress response (ESR) (4, 5). The ESR encompasses ~15% of the yeast genome, including ~600 repressed genes that are enriched for processes related to protein synthesis and growth, and the ~300 induced genes that encode diverse functions related to stress defense.

The discovery of a coordinated common response to stress suggested a possible mechanism for the long-observed phenomenon of acquired stress resistance and cross protection, where cells pretreated with a mild dose of stress are better able to survive an otherwise lethal dose of severe stress (6–9). In yeast, defective ESR expression correlates with diminished acquired stress resistance, suggesting that stress-activated gene expression changes may serve to protect against future challenges (10, 11). Beyond yeast, acquired stress resistance has been observed in diverse organisms including bacteria, plants, and humans, and has major implications for food production, agriculture, and human health. For example, mild stress induces higher resistance of the food-borne pathogens *Listeria monocytogenes* and *Salmonella typhimurium* to food preservatives (12, 13). Acquired stress resistance has also been implicated in host survival in the form of bile acid tolerance and antibiotic resistance (14, 15). Acquired thermotolerance and drought resistance in agriculturally significant plants is increasingly important due to climate change (16, 17). In humans, short-term fasting protects healthy cells, but not cancer cells, from the toxic effects of chemotherapy drugs (18, 19).Altogether, understanding how cells are able to acquire further resistance has broad applications ranging from agriculture and biotechnology to human health and disease.

We have been leveraging natural variation in the yeast ethanol response to better understand the cellular mechanisms governing acquisition to high stress resistance. While mutagenesis studies in laboratory strains of yeast have identified genes and processes necessary for acquired stress resistance (20–23), there are inherent limitations to using a single strain background (24). In the case of yeast, the S288c laboratory strain historically used for large-scale mutagenesis screening is a genetic and physiological outlier compared to wild yeast strains (25, 26). We have previously noted that S288c has an aberrant gene expression response to ethanol (11, 27), which we hypothesized was responsible for the strain’s inability to acquire resistance to any other stresses following pre-treatment with mild ethanol (10, 11). We subsequently found that defective induction of antioxidant defenses in response to mild ethanol was responsible for S288c’s inability to acquire further hydrogen peroxide (28).

In the present study we found that in contrast to S288c, mild ethanol stress induces cross protection against severe salt concentrations in the wild vineyard strain M22. This phenotype correlated with the induction of the Ena sodium efflux pump system by ethanol in a wild vineyard isolate, which was not induced in the S288c-derived common laboratory strain. Because the ENA system has been previously implicated in salt tolerance (29–31), we hypothesized that variation in *ENA* expression was responsible for phenotypic differences across strains. Surprisingly, we found that the *ENA* genes were completely dispensable for ethanol-induced survival of high salt concentrations in the wild strain. Instead, the *ENA* genes were necessary for a novel growth resumption phenotype that we call “ethanol-induced acclimation,” where mild ethanol stress allows cells to eventually resume growth on high concentrations of salt. More surprisingly, our common laboratory strain also exhibited ethanol-dependent acclimation to high salt concentrations, even though *ENA* is not induced by ethanol in this strain background. Overall, this study demonstrates that cross protection can affect both viability and growth through distinct mechanisms, both of which likely confer fitness effects that are ecologically relevant.

## RESULTS

### Ethanol Induces Cross Protection Against Severe Salt in a Wild Vineyard Strain

We previously observed that in our S288c-derived common laboratory strain, mild ethanol pretreatment could not induce acquired resistance to severe ethanol concentrations or cross protect against other stresses (10). In contrast, mild ethanol pretreatment did induce further ethanol resistance in the vast majority of wild yeast strains (11). Additionally, we recently discovered that mild ethanol stress can cross protect against oxidative stress in a wild oak strain (28). Because yeast in nature are likely to experience environmental shifts between high concentrations of sugars and high ethanol concentrations, we hypothesized that the inability of ethanol to cross protect against osmotic stress in S288c may be another aberrant acquired stress resistance phenotype in this strain background. We tested this by examining ethanol-induced cross protection against severe salt in a wild vineyard strain background (M22). Cross protection assays were performed by exposing cells to a mild dose of ethanol (5% v/v) for 1 hour, and scoring their survival across a panel of 11 increasingly severe doses of NaCl (see Materials and Methods).

We found that ethanol pretreatment weakly cross protected against severe NaCl in S288c (Fig. 1), though cross protection was not completely absent as previously reported (10). In contrast, ethanol strongly improved M22’s ability to survive otherwise lethal salt concentrations. Notably, S288c had intrinsically higher basal resistance to NaCl. However, the diminished cross protection phenotype of S288c relative to M22 cannot be explained by the higher baseline resistance, as mild NaCl pretreatment did strongly increase S288c’s NaCl resistance (Fig. 2). Moreover, the levels of acquired NaCl resistance following mild NaCl pretreatment were similar for both S288c and M22.

**Fig. 1.**
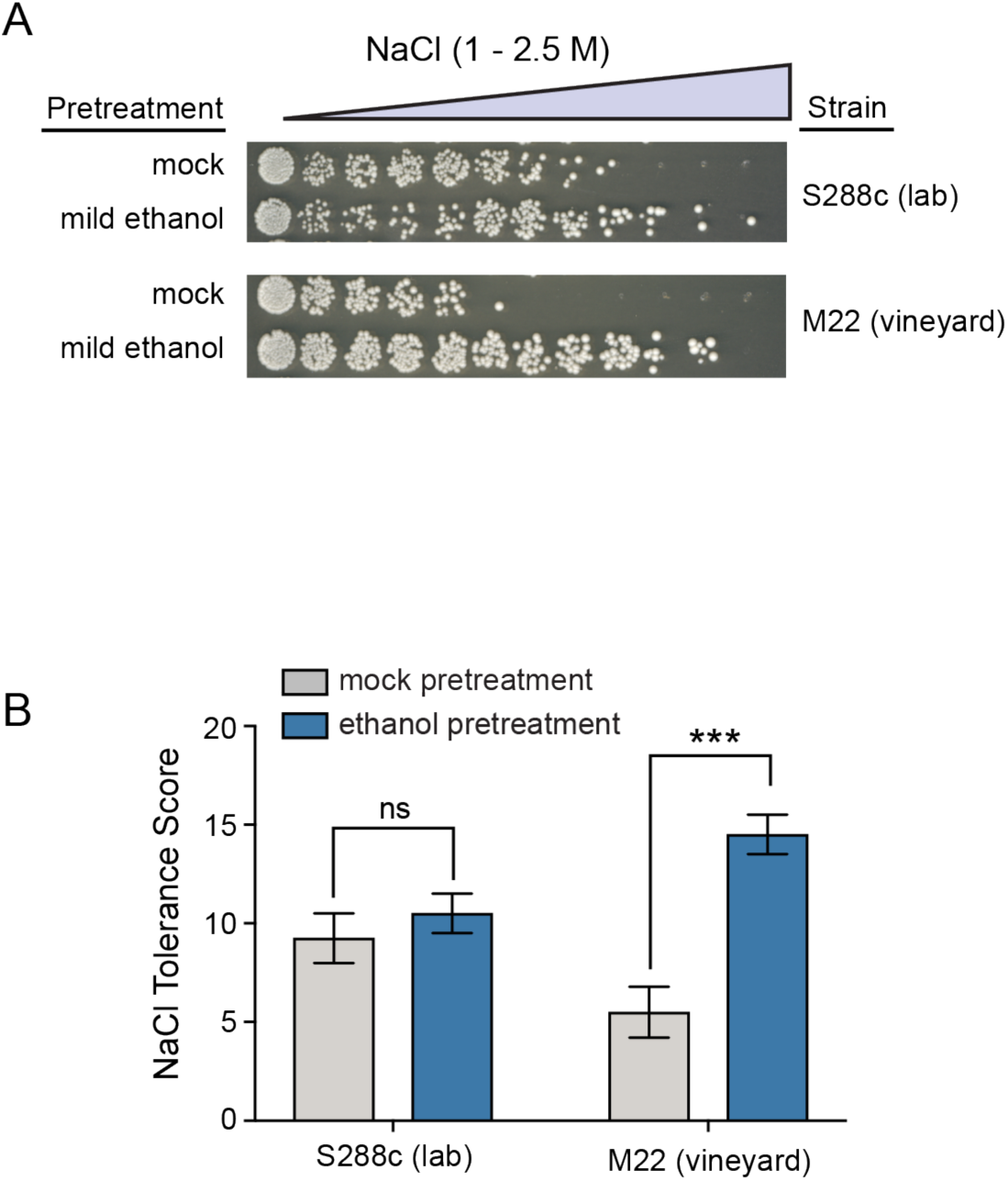
Ethanol induces cross protection against severe salt in a wild vineyard isolate. **A)** A representative NaCl cross protection assay. S288c (DBY8268 lab strain) and M22 (wild vineyard strain) were exposed to mild 5% ethanol or a mock (5% water) pretreatment for 1 hour, washed, and then exposed to 11 increasingly severe doses of NaCl for 2 hours before plating to score viability. **B)** NaCl tolerance scores were calculated from the viability across each of the 11 doses (see Methods). Error bars denote the standard deviation of four independent biological replicates. Asterisks indicate significantly higher resistance in ethanol-pretreated versus mock-pretreated cells (*** *P* < 0.001, ns = not significant (P > 0.05), *t*-test).

**Fig. 2.**
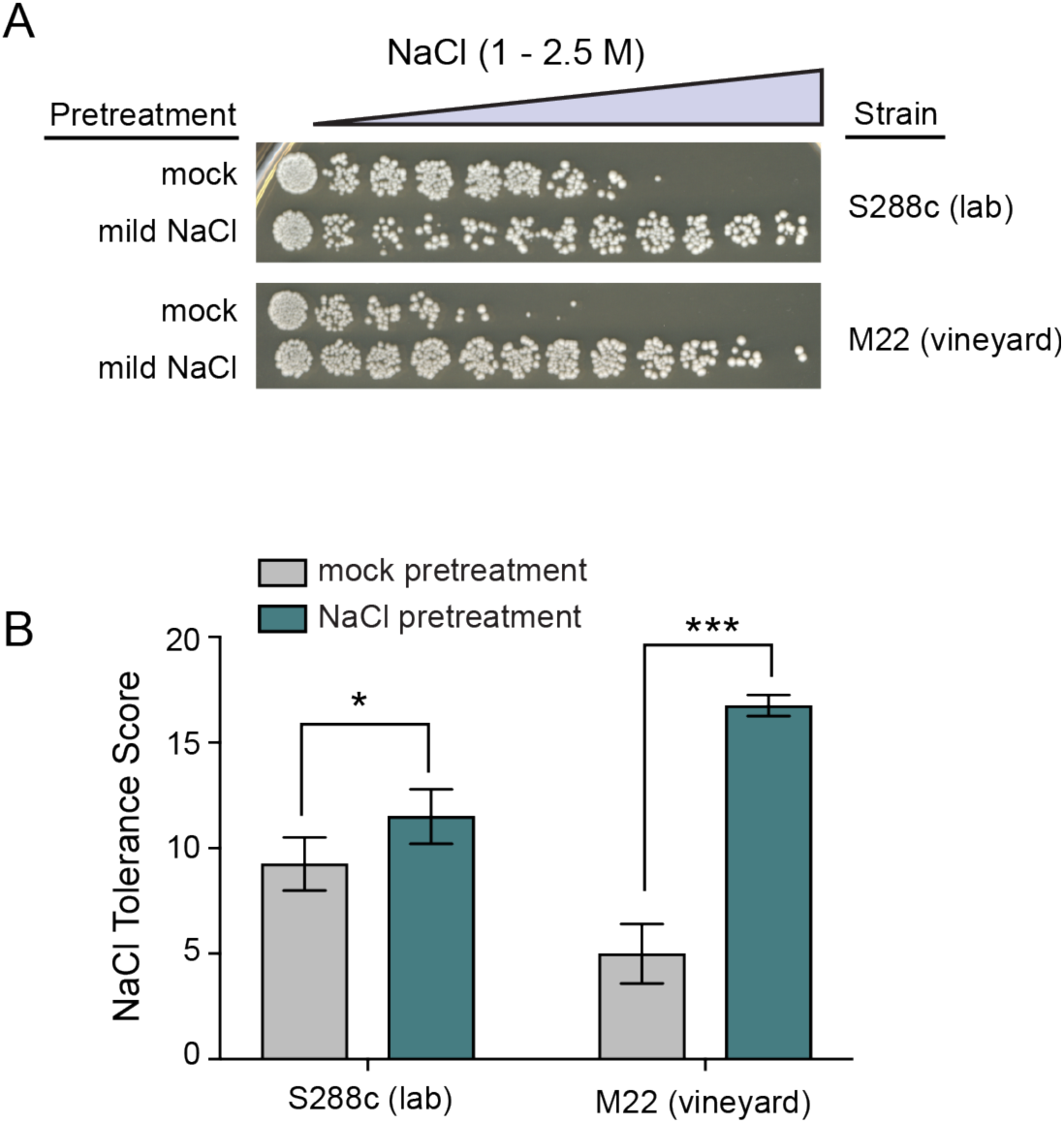
NaCl allows acquisition of even higher NaCl resistance in both the lab and vineyard strains. **A)** A representative acquired NaCl resistance assay is shown for both S288c (DBY8268 lab strain) and M22 (wild vineyard strain). Cells were split and exposed to either 0.4 M NaCl or a mock (media only) pretreatment for 1 hour, washed, exposed to 11 doses of severe NaCl for 2 hours, and then plated to score viability. **B)** Salt tolerance scores across each of the 11 doses are plotted as the mean and standard deviation of three independent biological replicates. Asterisks indicate significantly higher resistance in NaCl-pretreated versus mock-pretreated cells (* *P* < 0.05, *** *P* < 0.001, *t*-test).

### Induction of *ENA1* by Ethanol in a Wild Vineyard Isolate

Because acquired stress resistance relies on stress-activated gene expression changes (10, 11), we hypothesized that the phenotypic differences in cross protection between S288c and M22 may be due to differences in ethanol-responsive gene expression. We analyzed our previous ethanol-responsive transcriptome changes (27), specifically looking for candidate salt resistance genes with higher induction by ethanol treatment in M22 compared to S288c. We noticed that *ENA1* encoding a sodium efflux pump known to be involved in salt resistance (29) showed a 4.7-fold induction by ethanol in M22 versus a 1.4-fold decrease in expression in S288c, placing it within the top 25 genes in terms of magnitude of differential ethanol-responsive expression when comparing M22 and S288c.

The Ena P-type ATPase sodium efflux pumps are known to play a critical role in maintaining Na+ ion homeostasis in high salt conditions (30–32). In many yeast strain backgrounds, the *ENA* locus consists of a tandem array of nearly identical genes that can vary in copy number (33). S288c contains three copies *(ENA1, ENA2,* and *ENA5),* whereas M22 appears to contain a single copy (34). This single copy is somewhat unusual, in that M22 appears to contain a large 3885-bp deletion in *ENA1* relative to that of S288c, which results in a full-length in-frame fusion of *ENA1* and *ENA2* (34).

*ENA* copy number has been linked to high salt tolerance (25–27), likely explaining why S288c has higher basal NaCl tolerance than M22. In S288c, the *ENA* genes are lowly expressed under standard growth conditions (24), but are highly induced in response to saline or alkaline pH stresses (23). Including our previous studies, there are currently no reports of *ENA* induction by ethanol stress in the S288c, suggesting that this mode of *ENA* regulation may have been lost in the laboratory strain background (see Discussion).

### The ENA System is not Required for Ethanol-Induced Survival in Severe Salt, but is Required for Growth Resumption During Salt Acclimation

Because variation in *ENA* expression is linked to basal salt resistance, we hypothesized *ENA1* induction by ethanol may protect M22 from otherwise lethal salt stress. To test this, we deleted the entire *ENA* region in M22 and examined its role in ethanol-induced acquired NaCl resistance. Surprisingly, we found no defect in acquired salt resistance in the M22 *enaΔ* mutant (Fig. 3A). One possible explanation is that the ENA system is not necessary for short-term exposure to acute NaCl, but is instead required for long-term survival. Thus, we measured acquired NaCl resistance in the M22 *enaΔ* strain over 24 hours. Even after 24 hours, the M22 *enaΔ* strain acquired NaCl resistance equivalently to the wild-type strain, and had at most a mild basal NaCl resistance defect (Fig. 3A). We did notice that for ethanol pre-treated wild-type cells, the plating density significantly increased over time (Fig. 3B). This suggested that the ethanol pretreatment enabled cells to acclimate and resume growth on high concentrations of NaCl, and that this acclimation phenotype may be Ena dependent.

**Fig. 3.**
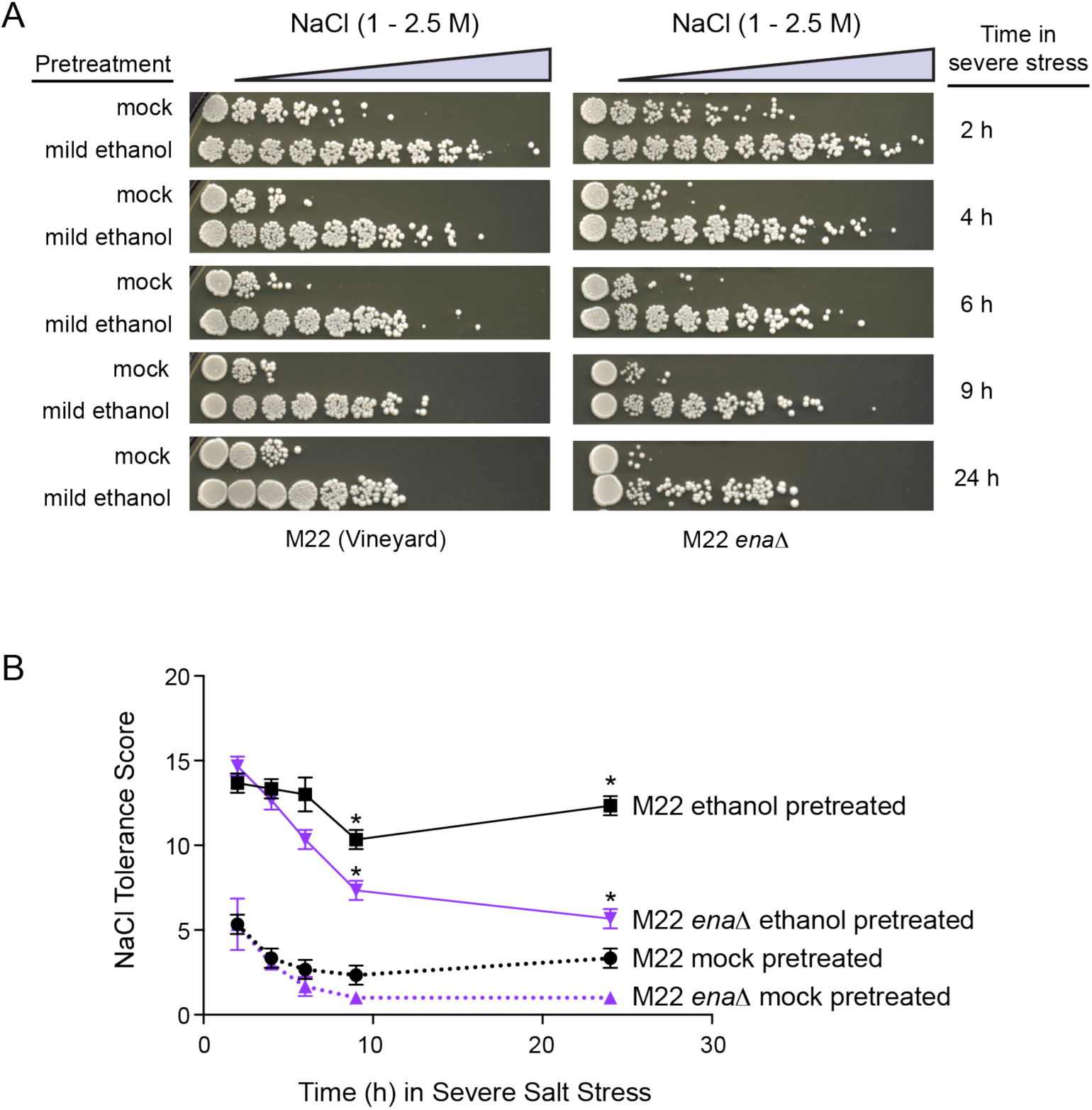
The ENA system is not necessary for ethanol-induced survival against severe salt. **A)** Representative NaCl cross protection assays are shown with increasing amounts of time in the severe secondary stress doses for Wild Type M22 and the M22 *enaΔ* mutant. **B)** Salt tolerance scores across each of the 11 doses were calculated for each of the timepoints from Panel A. Error bars denote the standard deviation of three independent biological replicates. Asterisks indicate timepoints with significantly higher resistance in M22 compared to the *enaΔ* mutant (*P < 0.01, *t*-test).

To further examine the role of the Ena in acclimation to high salt stress after ethanol pretreatment, we measured growth in liquid media, with or without mild ethanol pretreatment (Fig. 4A). Ethanol pretreatment allowed wild-type M22 cells to acclimate and resume growth in 1.25M NaCl, and this growth resumption was completely abolished in the M22 *enaΔ* mutant. These data suggest that two cross protection phenotypes—survival vs. growth—have distinct cellular mechanisms.

**Fig. 4.**
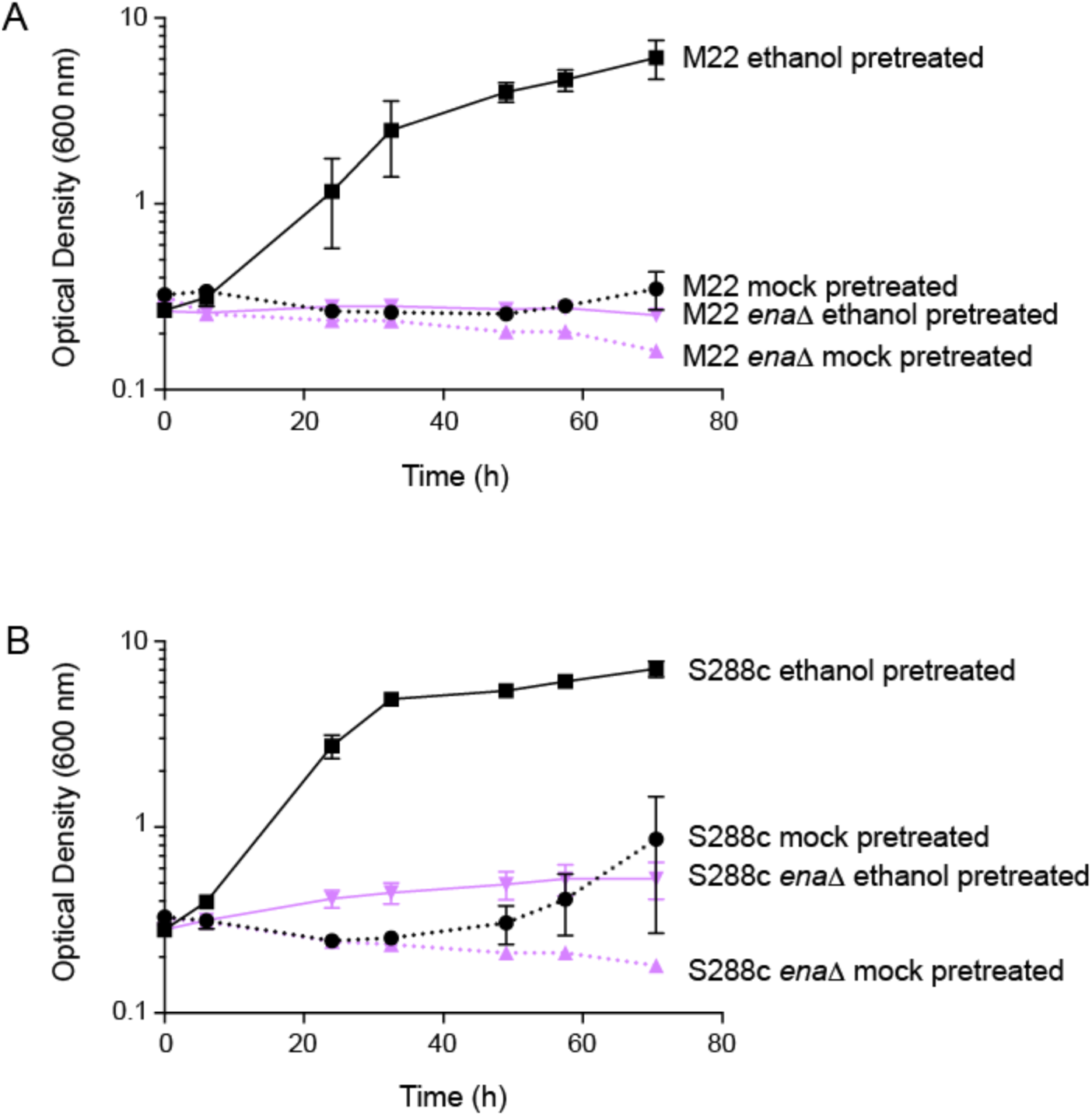
The ENA system is required for ethanol-induced growth resumption in severe salt in both S288c and M22. **A)** M22 and M22 *enaΔ* (JL213) and **B)** S288c (JL505) and S288c *enaΔ* (JL1131) were exposed to mild 5% ethanol or a mock (5% water) pretreatment for 1 hour, washed, and then resuspended in YPD containing 1.2 M NaCl. Growth was then measured over the course of 3 days. Error bars denote the standard deviation of three independent biological replicates.

In light of the distinct requirement of *ENA* for growth resumption on salt but not survival in the M22 background, we examined whether S288c was able to acclimate and resume growth on high salt following ethanol pretreatment. Indeed, ethanol pretreatment induced growth resumption in S288c (Fig. 4B), which was abolished in the S288c *enaΔ* mutant. This was somewhat surprising, considering that *ENA1* is not induced by ethanol in S288c under these conditions (see Discussion).

## DISCUSSION

In this study, we initially sought to understand how ethanol cross protects against severe salt stress. We found that ethanol-induced cross protection against severe salt was weaker in the common lab strain S288c when compared to the wild vineyard strain M22. We examined previous transcriptional profiling of the yeast ethanol response to identify candidate genes induced by ethanol in M22 but not S288c (27). Based on these data, we identified the ENA system as a prime candidate to test. The ENA system uses the hydrolysis of ATP through P-type ATPases to transport sodium out of the cell against the electrochemical gradient (22), and mutants lacking *ENA* function are salt sensitive (29–31). Interestingly, the *ENA* locus of many yeast strains including S288c and M22 appears to be the result of a recent introgression from *S. paradoxus* and shows significant copy number variation across strains (33). Other strains have a single non-S288c-like *ENA6* gene that does not share sequence similarity to the *ENA* genes from *S. paradoxus* (26, 30). Genetic mapping studies have linked both copy number variation and polymorphisms in the *ENA* region to variation in NaCl and LiCl tolerance (26, 35-37).

Thus, we were somewhat surprised to find that the ENA system of M22 was completely dispensable for survival in high salt following ethanol pretreatment. Instead, the ENA system was required for a novel ethanol-induced cross-protection phenotype that allows for acclimation and subsequent growth resumption in the presence of high salt. Notably, the vast majority of studies examining salt sensitivity phenotypes for *ENA* have been performed by growing cells on salt-containing plates, which cannot easily distinguish between viability and growth. Both phenotypes are likely important in natural environments. Wild yeast cells growing on fruit such as the M22 vineyard strain may experience simultaneous or fluctuating hyperosmotic stress and ethanol stress, which could explain the evolution of cross protection.

Because we were able to separately examine survival and growth, we reassessed the role of the ENA system in S288c. Surprisingly, we found that while ethanol only weakly induced higher survival on high salt in the S288c background, ethanol-induced acclimation to high salt was similar between the two strains. This ethanol-induced acclimation phenotype in S288c was also *ENA* dependent, despite the lack of induction of *ENA* by ethanol in this strain. Notably, *ENA* is known to be induced by NaCl in the S288c background (38), which is likely necessary for growth resumption on high salt. Additionally, basal *ENA* expression is higher in S288c compared to M22 (11, 27), likely due to copy number variation (our S288c-derived laboratory strain contains three *ENA* copies, while M22 contains a single copy (34)). It is likely that other ethanol-induced genes and processes are necessary for ethanol-induced acclimation to high salt concentrations.

The striking induction of the ENA system by ethanol in M22 but not S288c implies regulatory differences between the two strains. Recently, natural variation in the promoter region of *ENA6* in a sake strain was shown to increase Ena6p expression and thus increase salt tolerance (37). In this strain, a 33-bp deletion in the promoter eliminates glucose repression by eliminating repressor binding sites for the Nrg1p and Mig1/2p transcription factors. In contrast, we hypothesize that the novel regulation of *ENA1* by ethanol in M22 is likely not due to promoter variation. Comparing the promoters of *ENA1* between the S288c and M22 backgrounds reveals two SNPs and a 20-bp AT repeat insertion within a 20-bp poly-AT repeat region. However, these promoter differences do not alter or introduce any predicted transcription factor binding sites, suggesting promoter variation is unlikely responsible for the observed expression differences between the two strains. Instead, it is likely that *trans* regulatory variation is responsible for the novel induction by ethanol in the M22 background. The phenotypic consequences of this novel induction of *ENA1* by ethanol in the M22 strain remain an unresolved question, as S288c exhibits a similar growth resumption phenotype. Nonetheless, these findings expand our knowledge of the ENA system’s role in stress defense mechanisms, and highlight the power of using natural variation to yield new insight into even previously well-studied aspects of cellular physiology, such as the ENA system.

## MATERIALS AND METHODS

### Strains and Growth Conditions

Strains and primers used in this study are listed in Tables 1 and 2, respectively. The entire *ENA* region was deleted by homologous recombination and replacement with a KanMX4 drug resistance marker in the haploid MATa strain BY4741. This strain was used as a genomic template for introducing the *ena1-5Δ::KanMX4* allele into different strain backgrounds. To generate homozygous *enaΔ* diploids in the S288c strain background, the *ena1-5Δ::KanMX4* region was amplified and transformed into MATa and MATα haploid derivatives of DBY8268, which were then mated together. To generate homozygous *enaΔ* diploids in the M22 background, the *ena1-5Δ::KanMX4* region was transformed into the diploid M22 strain, resulting in an *enaΔ* heterozygote. M22 is capable of mating-type switching, and thus sporulation and dissection yielded homozygous *enaΔ* diploids. Homozygous deletions were verified by diagnostic PCR.

**Table 1.** Strains used in this study.

| Strain | Background | Genotype | Source |
| --- | --- | --- | --- |
| DBY8268 | S288c (lab strain) | *MATa/MATα ura3-52/ura3Δ ho/ho GAL2/GAL2* | David Botstein |
| JL187 | BY4741 | *MATa his3Δ leu2Δ met15Δ ura3Δ ena1-5Δ::kanMX* | This study |
| JL505 | DBY8268 | *MATa ho GAL2 ura3-52 or ura3Δ* |  |
| JL506 | DBY8268 | *MATα ho GAL2 ura3-52 or ura3Δ* |  |
| JL1127 | DBY8268 | *MATa ho GAL2 ura3-52 or ura3Δ ena1-5Δ::KanMX* | This study |
| JL1128 | DBY8268 | *MATα ho GAL2 ura3-52 or ura3Δ ena1-5Δ::KanMX* | This study |
| JL1131 | DBY8268 | *MATa/MATα ho/ho GAL2/GAL2 ura3-52/ura3Δ ena1-5Δ::KanMX/ena1-5Δ::KanMX* | This study |
| M22 | wild vineyard strain | *MATa/MATα* | Robert Mortimer |
| JL213 | M22 | *MATa/MATα ena1-5Δ::KanMX/ena1-5Δ::KanMX* | This study |
| JL214 | M22 | *MATa/MATα ena1-5Δ::KanMX/ENA1-5* | This study |

**Table 2.**
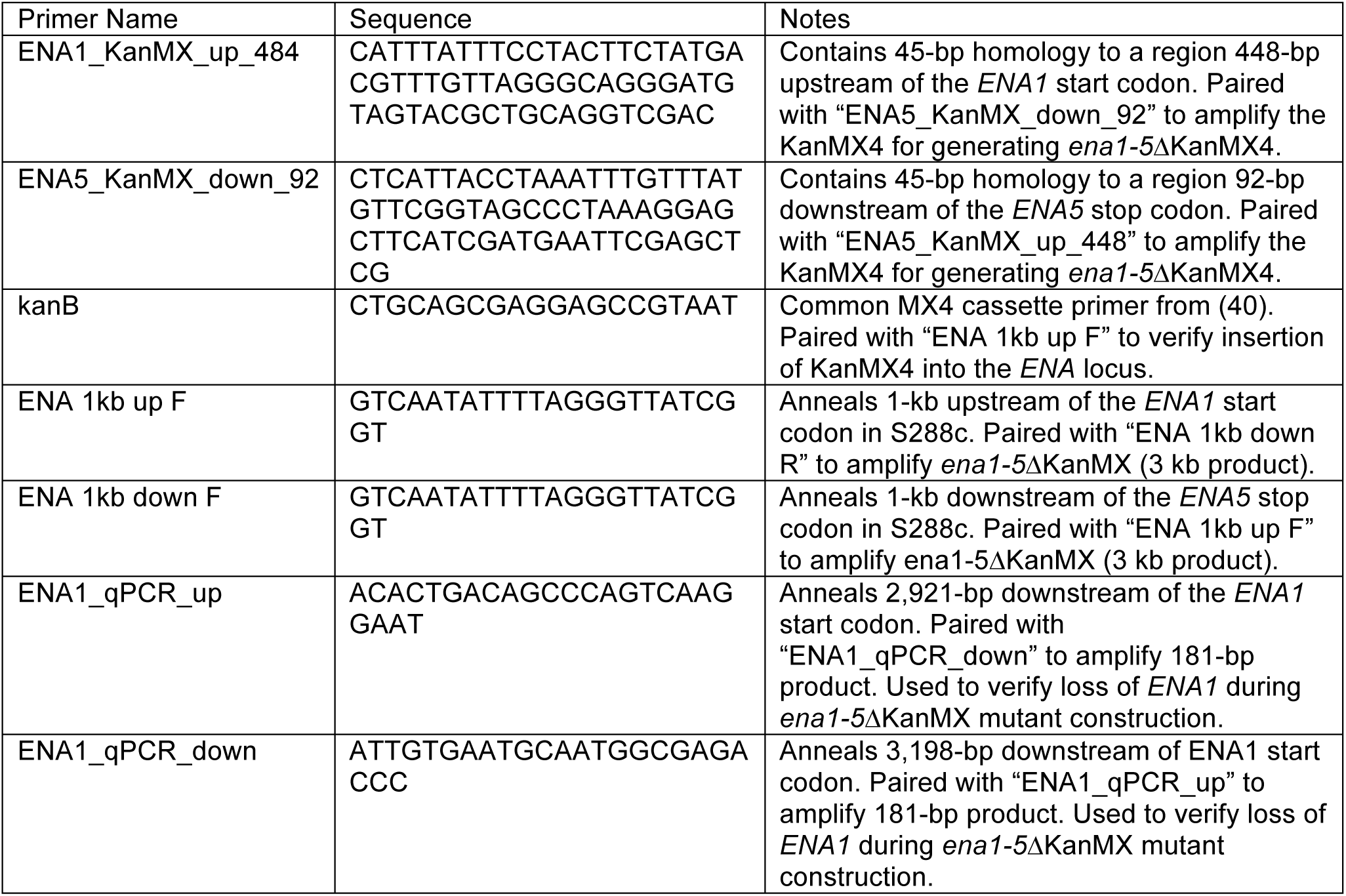
Primers used in this study.

All strains were grown in YPD (1% yeast extract, 2% peptone, 2% dextrose) at 30°C with orbital shaking (270 rpm). Optical density was recorded using a Unico spectrophotometer. Sporulation was achieved by growing cells to saturation for 2 days in YPD, harvesting by centrifugation, resuspending in 1% potassium acetate, and incubating for up 3-5 days at 25°C with shaking.

### Acquired Stress Resistance Assays

The acquired stress resistance assays were performed as described in (28). Briefly, cells were grown to overnight to saturation, sub-cultured into in 30-mL fresh media, and then grown for at least 8 generations into exponential phase (OD_600_ of 0.3-0.6) to reset any cellular memory of starvation stress (39). Each culture was split and pretreated with either a mild “primary” stress or a mock (equivalent concentration of water) control. Primary stresses included either 5% v/v ethanol or 0.4 M NaCl. Cells were incubated with the pretreatment for 1 hour and then collected by mild centrifugation at 1,500 *x g* for 3 min to remove the primary stress. Cells were resuspended in fresh media to an OD_600_ of 0.6, and then diluted 3-fold into a microtiter plate containing a panel of severe NaCl doses ranging from 1.2 M to 3.2 M (0.2 M increments). Plates were sealed breathable Rayon films (VWR), and incubated with secondary stress at 30°C with 800 rpm shaking in a VWR Symphony Incubating Microplate Shaker. Secondary treatments were for 2h unless otherwise noted. Following secondary treatment, 4 μl of a 50-fold cell dilution was spotted directly onto YPD agar plates and grown for 48 hours at 30°C. Viability at each dose was scored using a 4-point semi-quantitative scale that compared survival in each secondary dose against an unstressed (YPD only) control: 100% viability = 3 points, 50-90% viability = 2 points, 10-50% viability = 1 point, and 0% viability = 0 points. An overall tolerance score was calculated as the sum of scores across all 11 stress doses. Acquired stress resistance assays were performed in biological triplicate, and raw phenotypic data can be found in Table S3. A detailed acquired stress resistance assay protocol can be found on protocols.io under doi dx.doi.org/10.17504/protocols.io.g7sbzne. Statistical analyses were performed using Prism 7 (GraphPad Software).

### Ethanol-Induced Growth Resumption Analysis

To assess ethanol-induced growth resumption in the presence of salt, cells were given mild primary ethanol (5% v/v) or mock pretreatments as described for the acquired resistance assays. Following 1 hour pretreatment, cells were gently centrifuged at room temperature for 3 minutes at 1500 *x g*, and then resuspended in YPD containing 1.25 M NaCl at an OD_600_ of 0.1. Five ml of each sample was transferred to a glass test tube and incubated at 30°C at 270 rpm. The OD_600_ of all samples was then manually measured with a Unico spectrophotometer over approximately 72 hours. Growth assays were performed in biological triplicate.

## ACKNOWLEDGEMENTS

This work was supported in part by National Science Foundation Grant No. I0S-1656602, the Arkansas Biosciences Institute (Arkansas Settlement Proceeds Act of 2000), and startup funds provided by the University of Arkansas to JAL, University of Arkansas Honors College Grants to EAM and MV, and the DOE Great Lakes Bioenergy Research Center (DOE Office of Science BER DE-FC02-07ER64494) The funders had no role in study design, data collection and interpretation, or the decision to submit the work for publication.

